# Widespread Distribution of α-Synuclein Oligomers in *LRRK2*-related Parkinson’s Disease

**DOI:** 10.1101/2024.12.18.629265

**Authors:** Hiroaki Sekiya, Lukas Franke, Yuki Hashimoto, Mariko Takata, Katsuya Nishida, Naonobu Futamura, Kazuko Hasegawa, Hisatomo Kowa, Owen A. Ross, Pamela J. McLean, Tatsushi Toda, Zbigniew K. Wszolek, Dennis W. Dickson

## Abstract

Mutations in leucine-rich repeat kinase 2 (*LRRK2*) are the most common cause of familial and sporadic Parkinson’s disease (PD). While the clinical features of *LRRK2*-PD patients resemble those of typical PD, there are significant differences in the pathological findings. The pathological hallmark of definite PD is the presence of α-synuclein (αSYN)-positive Lewy-related pathology; however, approximately half of *LRRK2*-PD cases do not have Lewy-related pathology. Lewy-related pathology is a late-stage αSYN aggregation that can be visualized with hematoxylin and eosin stains or conventional immunohistochemistry (IHC). Increasing evidence has indicated that αSYN oligomers, which represent the early-stage of αSYN aggregation, may have neurotoxicity. Visualization of αSYN oligomers requires specialized staining techniques, such as αSYN-proximity ligation assay (PLA). The distribution and severity of αSYN oligomers in the human brain of *LRRK2*-PD patients remain unknown. In this study, we performed phosphorylated αSYN-IHC and αSYN-PLA staining on postmortem brain sections of patients with three pathogenic LRRK2 mutants: p.G2019S (n=5), p.I2020T (n=5), and p.R1441C (n=4). The severity of Lewy-related pathology and αSYN oligomers were assessed semi-quantitatively in the brainstem, limbic lobe, basal ganglia, and cerebral cortex. αSYN oligomers were detected in *LRRK2*-PD cases even in cases without Lewy-related pathology; a negative correlation was observed between Lewy-related pathology and αSYN oligomers (r=-0.26 [-0.39, -0.12]; P<0.0001). Our findings suggest that αSYN oligomers may represent a common pathological feature of *LRRK2*-PD. Notably, patients harboring p.G2019S and p.I2020T had significantly higher levels of αSYN oligomers in those without Lewy-related pathology compared to those with Lewy-related pathology. These cases also had a trend toward shorter disease duration. These results imply that in *LRRK2*-PD, αSYN oligomers may initially accumulate in the brain but do not progress to form Lewy-related pathology. The present study suggests that targeting αSYN oligomers may be a therapeutic strategy for *LRRK2*-PD even if there is no Lewy-related pathology.

## Introduction

Parkinson’s disease (PD) is the second most common neurodegenerative disorder, clinically characterized by bradykinesia, tremor, and rigidity [34]. The pathological hallmark of PD is the presence of α-synuclein (αSYN)-positive intracellular inclusions, called Lewy bodies, which are found widespread in selectively vulnerable neurons, as well as neuronal loss in the substantia nigra [11, 12, 26]. While most PD cases are considered sporadic, relatively rare familial forms of PD are widely recognized, and genetic factors also play an important role in sporadic PD [5, 6, 16]. Mutations in the *LRRK2* gene, which encodes the leucine-rich repeat kinase 2 (LRRK2) protein, were identified in a group of familial PD patients [14, 15, 33, 55]. Mutations in *LRRK2* represent the most common genetic cause of PD [13, 20, 45]. LRRK2 is a large multidomain protein with two primary enzymatic domains: a GTPase domain and a kinase domain [3, 32, 37]. The most frequent pathogenic substitutions are found in these enzymatic domains, such as p.R1441C/G/H and Y1669C in the GTPase domain and p.G2019S and p.I2020T in the kinase domain [17, 24, 35, 50]. While it is reported that disease progression is slightly slower in *LRRK2*-PD than in sporadic PD [41], the clinical manifestations of *LRRK2*-PD patients are largely indistinguishable from those of typical PD [19, 46, 47]. In contrast, the neuropathology of *LRRK2*-PD is heterogeneous, with inconsistent presence of Lewy bodies even within the same family. Kalia et al. reported that 65% (11/17) of p.G2019S, 11% (1/9) of p.I2020T, and 50% (2/4) of p.R1441C carriers had Lewy bodies [25], while Henderson et al. reported that only 56% (5/9) of p.G2019S patients had Lewy bodies [21].

While Lewy bodies are late-stage aggregates of αSYN, αSYN oligomers represent an earlier stage of αSYN aggregation. Increasing evidence indicates that αSYN oligomers may be more toxic than Lewy bodies [8, 51]. Lewy bodies are usually round inclusions in the neuronal cytoplasm with hematoxylin-eosin stains, while they are more numerous and associated with extraneuronal pathology with immunohistochemistry for phosphorylated αSYN (p-αSYN). In contrast, αSYN oligomers are not visible with routine histologic methods or even with p-αSYN immunohistochemistry. Oligomer detection requires special staining methods. Recently, the proximity ligation assay (PLA) has been developed to visualize αSYN oligomers in pathological specimens [36, 43, 44, 54]. Our previous work showed a widespread distribution of αSYN oligomers in PD [44]; however, to date, the distribution of αSYN oligomers in PD with *LRRK2* mutations has not yet been examined. Given the similarities in clinical manifestations of sporadic PD and PD with *LRRK2* mutations, we hypothesized that αSYN oligomers would be present in *LRRK2*-PD. The present study investigated the distribution of αSYN oligomers in *LRRK2*-PD with and without Lewy-related pathology.

## Materials and methods

### Subjects

The present study examined PD patients with three different *LRRK2* mutations: p.G2019S, p.I2020T, and p.R1441C. Post-mortem brain samples were obtained from the National Hospital Organization Sagamihara Hospital (Sagamihara, Kanagawa, Japan), the National Hospital Organization Hyogo Chuo National Hospital (Sanda, Hyogo, Japan), and the Mayo Clinic brain bank (Jacksonville, Florida, USA). This study included five patients with p.G2019S, five with p.I2020T, four with p.R1441C, and five control subjects. A subset of patients with p.G2019 [39], p.I2020T [18, 48], and p.R1441C [52, 53, 55] was previously described. The control subjects did not have parkinsonism and they had no Lewy-related pathology with phosphorylated αSYN immunohistochemistry (IHC).

### Brain samples

Postmortem brain samples were processed as previously described [18, 42]. Each brain was fixed in 10% formalin and representative brain regions were embedded in paraffin after fixation. Serial sections (6-µm) were stained with hematoxylin and eosin for neuropathological evaluation. The following regions were included: dorsal motor nucleus of the vagus, medullary raphe nuclei, locus coeruleus, substantia nigra, raphe nuclei of the midbrain, entorhinal cortex, hippocampus, putamen, caudate nucleus, and neocortex (frontal, temporal, parietal, and occipital). Two contiguous sections of each region were stained: one with phosphorylated-αSYN immunohistochemistry and one with αSYN-PLA.

All brain autopsies were performed with the consent of the legal next-of-kin or an individual with legal authority to grant permission for autopsy. The study was approved by the Kobe University Ethical Committee (2010-073). In addition, de-identified studies using autopsy samples are considered exempt from human subject research by the Mayo Clinic Institutional Review Board.

### Staining protocols

Lewy-related pathology was visualized using IHC with an antibody for phosphorylated-αSYN (1:2000; mouse monoclonal, psyn#64, FUJIFILM Wako Pure Chemical Corporation, Osaka, Japan) as previously described [43, 44].

To visualize αSYN oligomers, we conducted αSYN-PLA staining using Duolink kits supplied by Sigma-Aldrich (St. Louis, MO, USA) as previously described [43, 44]. PLA probes were conjugated to anti-αSYN antibodies (mouse monoclonal, Syn211, Abcam, Cambridge, UK). Twenty micrograms of Syn211 antibody were added to 2 µl of conjugation buffer and the solution was transferred to a vial of lyophilized oligonucleotides. After incubating the solution at room temperature overnight, 2 µl of stop solution was added and incubated for 30 min at room temperature. This solution was suspended in 24 µl of storage solution. Paraffin-embedded brain sections were dewaxed in xylene and rehydrated in a graded series of alcohol. After antigen retrieval and blocking, the tissues were incubated with PLA probe conjugated Syn211 antibodies at 37°C for 1 hour, and then at 4°C overnight. After washing the sections with wash buffer and blocking endogenous peroxidase, ligation was performed at 37°C for 1 hour, followed by amplification at 37°C for 2.5 hours. The sections were then incubated with detection solution at room temperature for 1 hour and finally with substrate solution at room temperature for 20 minutes. Subsequently, the sections were counterstained with hematoxylin, dehydrated through graded alcohols and xylene, and coverslipped with mounting media.

### Evaluation of each pathology

The severity of Lewy-related pathology and αSYN oligomers was assessed as previously described [44]. We assessed the Lewy-related pathology burden on phosphorylated-αSYN IHC slides using a five-point scale (0: absent, 1: slight, 2: mild, 3: moderate, and 4: severe) at 20x magnification. We similarly assessed the severity of αSYN oligomers on αSYN-PLA stained slides at 20x magnification. Neuronal staining patterns and neuropil staining severity were assessed. Neuronal staining was classified based on the staining pattern as follows: neuronal-clustered, neuronal-patchy, neuronal-punctate, and null (no signal detected) (**online resource Fig. S1A**). Neuronal-clustered was defined as αSYN-PLA signals throughout the neuronal perikarya; neuronal-patchy had patchy αSYN-PLA signals in the neurons; and neuronal-punctate had dot-like αSYN-PLA signals. For comparison, neuronal staining patterns were converted to numerical values. Specifically, neuronal-cluster was converted to 3, neuronal-patchy to 2, neuronal-punctate to 1, and null to 0. Subsequently, neuropil staining was scored on a six-point scale (0: no to 5: highest) by using pre-made scoring plates (**online resource Fig. S1B**). For the cortical sections, we eliminated bias by evaluating the severity of αSYN-PLA staining in neuropil in the three most affected microscopic fields and calculating the average scores. In other regions, we used areas with the most abundant signals. Then, to compare the severity of αSYN oligomer burden with Lewy-related pathology burden, we made a combined severity score for αSYN oligomers on a five-point scale from the neuronal staining pattern and neuropil scores.

In addition to the semi-quantitative scoring, a quantitative measure of the stained area of αSYN-PLA staining was performed using the software ImageJ (National Institute of Health, Bethesda, MD, USA) as previously described [44]. The cortical images were used because they were large enough to fill all areas of 1720 x 1075 pixels at 20x magnification. Three representative images of each cortex were obtained with the Aperio AT2 Slide Scanner (Leica Biosystems, Deer Park, IL, USA) at 20x magnification. Quantification of the stained area was performed using the IHC Image Analysis Toolbox plugin. After the training procedure with Nova Red color, the stained images were analyzed with color detection and converted to a 16-bit format (**online resource Fig. S2A**) for measurement of the area of pixels surpassing the threshold.

### Statistical analysis

All statistical analyses were conducted by using GraphPad Prism (version 9.1.2, GraphPad Software, La Jolla, CA, USA). Fisher’s exact test was used to compare categorical variables. For continuous variables, an unpaired t test was used. The Mann-Whitney test was conducted to compare pathological severity scores. Spearman correlation was used to assess the relationship between pathological severity scores and stained area. For the comparison of αSYN oligomer scores in each brain region between *LRRK2* mutation carriers and control subjects, the Kruskal-Wallis test was conducted. As a post-hoc analysis, Dunn’s multiple comparison test was performed between each mutation and the control group. Statistical significance was defined by a p-value < 0.05.

## Results

### Patient characteristics

**Table 1** summarizes the demographic and clinical features of patients and subjects. There was no significant difference in age at death between patients with each mutation and controls. Disease duration was comparable among patients with each mutation. **Table 2** summarizes the clinical information of each patient with *LRRK2*-PD. Initial symptoms were gait difficulty in 2 and tremor in 3 out of 5 p.G2019S patients, gait difficulty in 3, rigidity in 1, and bradykinesia and rigidity in 1 out of 5 p.I2020T patients, and bradykinesia in 2 and tremor in 2 out of 4 p.R1441C patients.

**Table 1.**
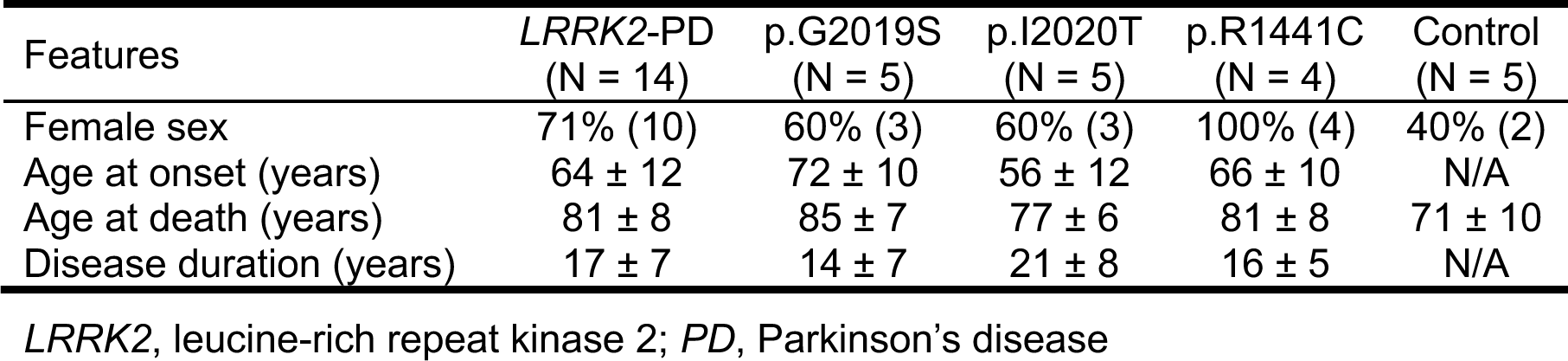
Demographic and clinical features.

**Table 2.** Clinical information of LRRK2-PD patients.

| ID | Mutations | Sex | AAO (y) | AAD (y) | Disease duration (y) | Initial symptoms |
| --- | --- | --- | --- | --- | --- | --- |
| L1 | p.G2019S | F | 73 | 80 | 6.1 | Gait difficulty, falls |
| L2 | p.G2019S | F | 82 | 89 | 6.9 | Gait difficulty |
| L3 | p.G2019S | M | 55 | 77 | 22.2 | Tremor |
| L4 | p.G2019S | M | 68 | 84 | 16.2 | Tremor |
| L5 | p.G2019S | F | 77 | 94 | 16.6 | Tremor |
| L6 | p.I2020T | F | 49 | 78 | 28.8 | Rigidity |
| L7 | p.I2020T | F | 70 | 80 | 10.8 | Gait difficulty |
| L8 | p.I2020T | M | 56 | 81 | 24.3 | Gait difficulty |
| L9 | p.I2020T | M | 39 | 67 | 28.1 | Gait difficulty |
| L10 | p.I2020T | F | 63 | 77 | 13.9 | Bradykinesia, rigidity |
| L11 | p.R1441C | F | 57 | 68 | 11.2 | Bradykinesia |
| L12 | p.R1441C | F | 59 | 79 | 20.0 | Tremor |
| L13 | p.R1441C | F | 78 | 89 | 11.6 | Bradykinesia |
| L14 | p.R1441C | F | 68 | 88 | 20.8 | Tremor |

*ID*, patient number; *AAO*, age at onset; *AAD*, age at death; *y*, years

Lewy-related pathology was observed in 3 out of 5 p.G2019S, 2 out of 5 p.I2020T, and 1 out of 4 p.R1441C patients. Clinical features were compared between patients with and without Lewy-related pathology (**Table 3**). Disease duration tended to be longer in patients with Lewy-related pathology compared to those without Lewy-related pathology (20 vs. 15 years, P=0.25). Ages at onset and death were comparable between patients with and without Lewy-related pathology (P=0.58 and P=0.86).

**Table 3.** Features between patients with and without Lewy-related pathology.

| Feature | Lewy-related<br>pathology (+)<br>(N = 6) | Lewy-related<br>pathology (-)<br>(N = 8) | P<br>value |
| --- | --- | --- | --- |
| Female sex | 50% (3) | 88% (7) | 0.24 |
| Age at onset | 62 ± 13 | 66 ± 12 | 0.58 |
| Age at death | 82 ± 10 | 81 ± 7 | 0.86 |
| Disease duration | 20 ± 5 | 15 ± 8 | 0.25 |
| Mutations |  |  |  |
| p.G2019S | 60% (3) | 40% (2) | - |
| p.I2020T | 40% (2) | 60% (3) | - |
| p.R1441C | 25% (1) | 75% (3) | - |

### Distribution and severity of αSYN oligomers

The distribution and severity of αSYN oligomers were assessed on αSYN-PLA stained slides. Representative images of αSYN-staining are shown in **Figure 1**. The left images depict patients who had no Lewy-related pathology, while the right images represent patients who had Lewy-related pathology. Since signals were observed in both neuropil and neurons, the severity for each was evaluated separately, as previously described [43, 44]. To ensure objectivity of neuropil scoring, we examined the correlations with stained area. There was a significantly strong correlation between the neuropil score and stained area (r=0.72 [0.59, 0.81]; P<0.0001, **online resource Fig. S2B**). The combined score, which was generated based both on neuropil score and neuronal staining patterns, also showed a significant correlation with the stained area (r=0.73 [0.61, 0.82]; P<0.0001, **online resource Fig. S2C**).

**Figure 1.**
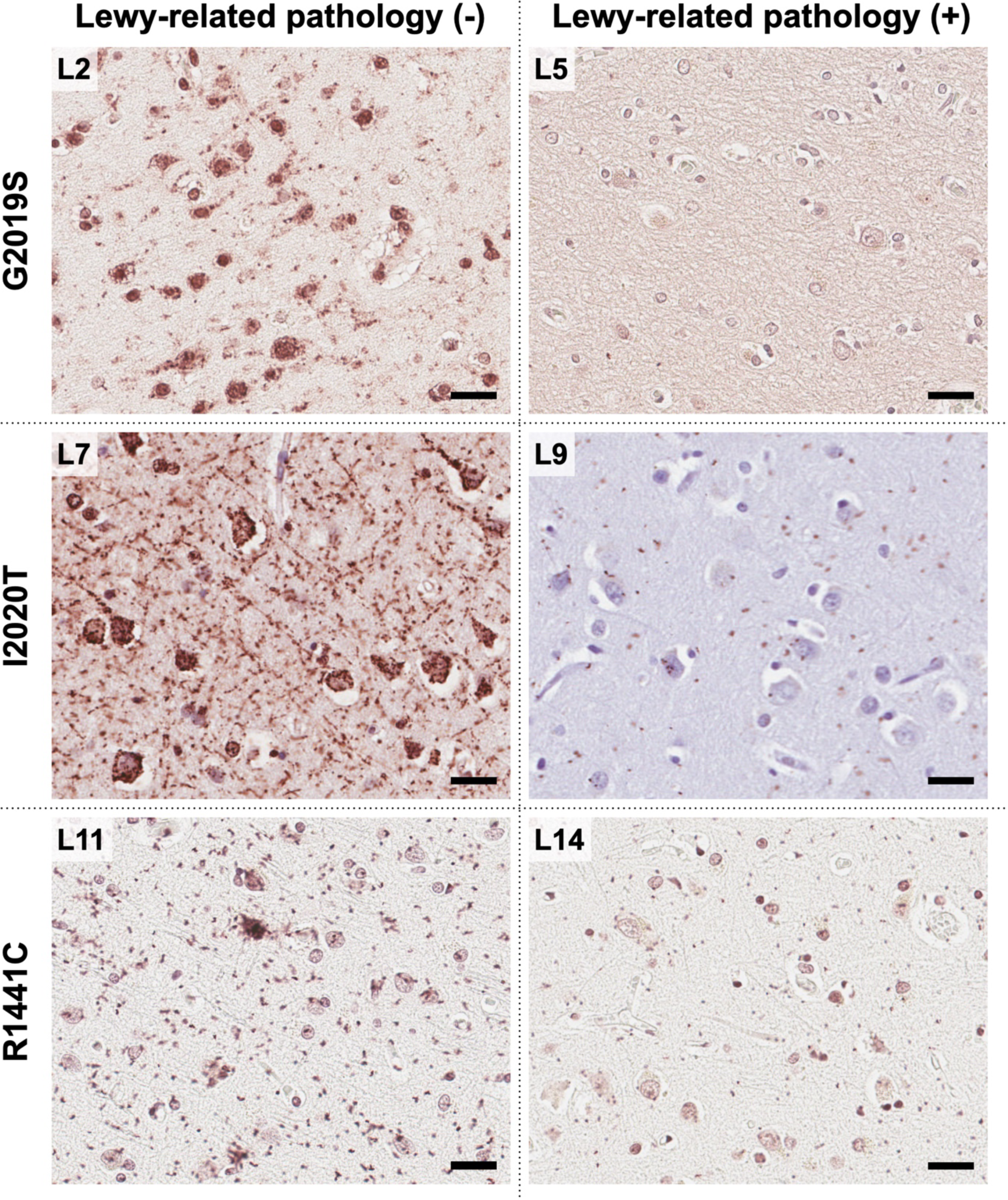
Representative images of αSYN-PLA staining. Left images are from patients without Lewy-related pathology and right images are from patients with Lewy-related pathology. The red-brown signals indicate αSYN oligomers. These αSYN oligomers accumulate in neurons and neuropil. Cases without Lewy-related pathology exhibit more abundant αSYN oligomers compared to those with Lewy-related pathology. Scale bar: 50 µm.

First, we assessed the staining patterns of neurons as neuronal-clustered, neuronal patchy, neuronal-punctate, or null. **Figure 2A** summarizes the results. The neuronal-clustered pattern was found in one of the p.G2019S patients (L2) and two of the p.I2020T patients (L6 and L7). All patients with neuronal-clustered staining pattern lacked Lewy-related pathology. Most brain regions in control subjects did not have any staining in neurons. Each staining pattern was converted into a numerical value for comparison. *LRRK2*-PD patients exhibited a more severe burden in neurons compared to control subjects (P<0.0001). More severe staining was observed in patients without Lewy-related pathology compared to those with Lewy-related pathology in p.G2019S and p.I2020T (P<0.0001 and P=0.0017), while no difference was observed in p.R1441C (P>0.99).

**Figure 2.**
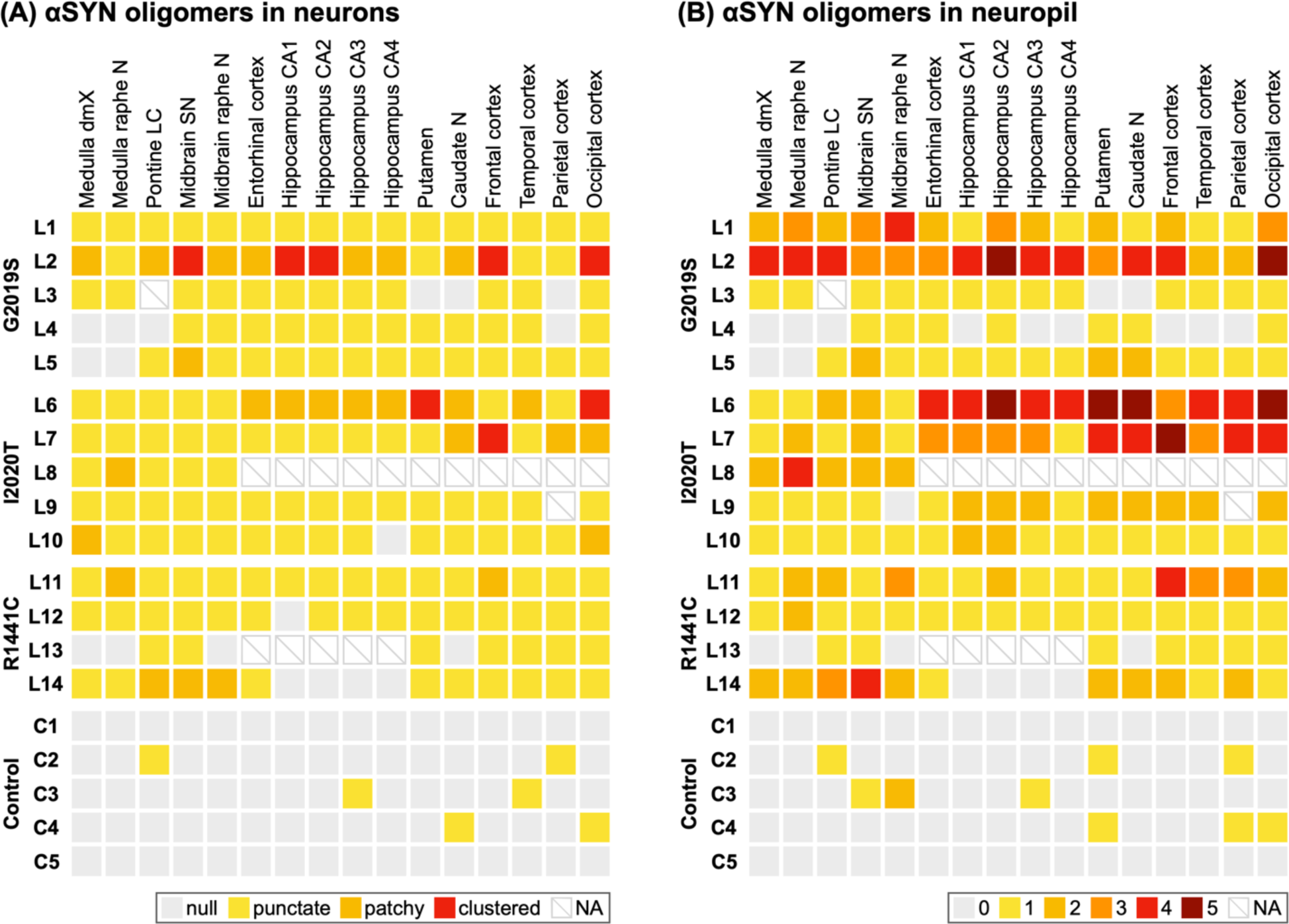
Neuronal staining patterns (A) and the severity in neuropil (B) of αSYN-PLA staining for each brain region. The staining pattern of αSYN oligomers in neurons shows a clustered appearance in one patient with p.G2019S and two patients with p.I2020T mutations. Regarding αSYN oligomers in the neuropil, two patients of each mutation exhibit severity scores of 4 or 5. *αSYN*, α-synuclein; *PLA*, proximity ligation assay; *dmX*, dorsal motor nucleus of the vagal nerve; *N*, nucleus; *LC*, locus coeruleus; *SN*, substantia nigra; *CA*, cornu ammonis.

Next, we assessed the severity of neuropil staining (**Figure 2B**). The most severe burden, with scores of 4 or 5, was observed in two p.G2019S patients (L1 and L2), three p.I2020T patients (L6, L7, and L8), and two p.R1441C patients (L11 and L14). All cases with severe burdens lacked Lewy-related pathology, except for one p.R1441C case. Within each mutation, αSYN oligomer burden in neuropil was greater in patients without Lewy-related pathology than in those with Lewy-related pathology for p.G2019S and p.I2020T (P<0.0001 and P=0.0002). For p.R1441C, there was no difference in neuropil αSYN oligomer burden between patients with and without Lewy-related pathology (P=0.53). Most brain regions in control subjects did not show αSYN oligomers in the neuropil. As a whole, *LRRK2*-PD patients had higher neuropil scores compared to controls (P<0.0001).

### Comparison of αSYN oligomers and Lewy-related pathology

To understand the overall αSYN oligomer burden and compare it to the burden of Lewy-related pathology, a combined αSYN oligomer score was generated from the neuropil score and neuronal staining patterns. The severity of αSYN oligomers and Lewy-related pathology in each brain region is summarized in **Figure 3**. The highest oligomer burden was found in one case of p.G2019S and two cases of p.I2020T, both without Lewy-related pathology. There was a negative correlation between burdens of αSYN oligomer and Lewy-related pathology (r=-0.26 [-0.39, -0.12]; P<0.0001; **online resource Fig. S3**). Within each mutation, the combined αSYN oligomer score was higher in patients without Lewy-related pathology than that in patients without Lewy-related pathology for p.G2019S and p.I2020T (P<0.0001 and P<0.0001). In patients with p.R1441C, there was no significant difference in combined αSYN oligomer score between patients with and without Lewy-related pathology (P=0.66).

**Figure 3.**
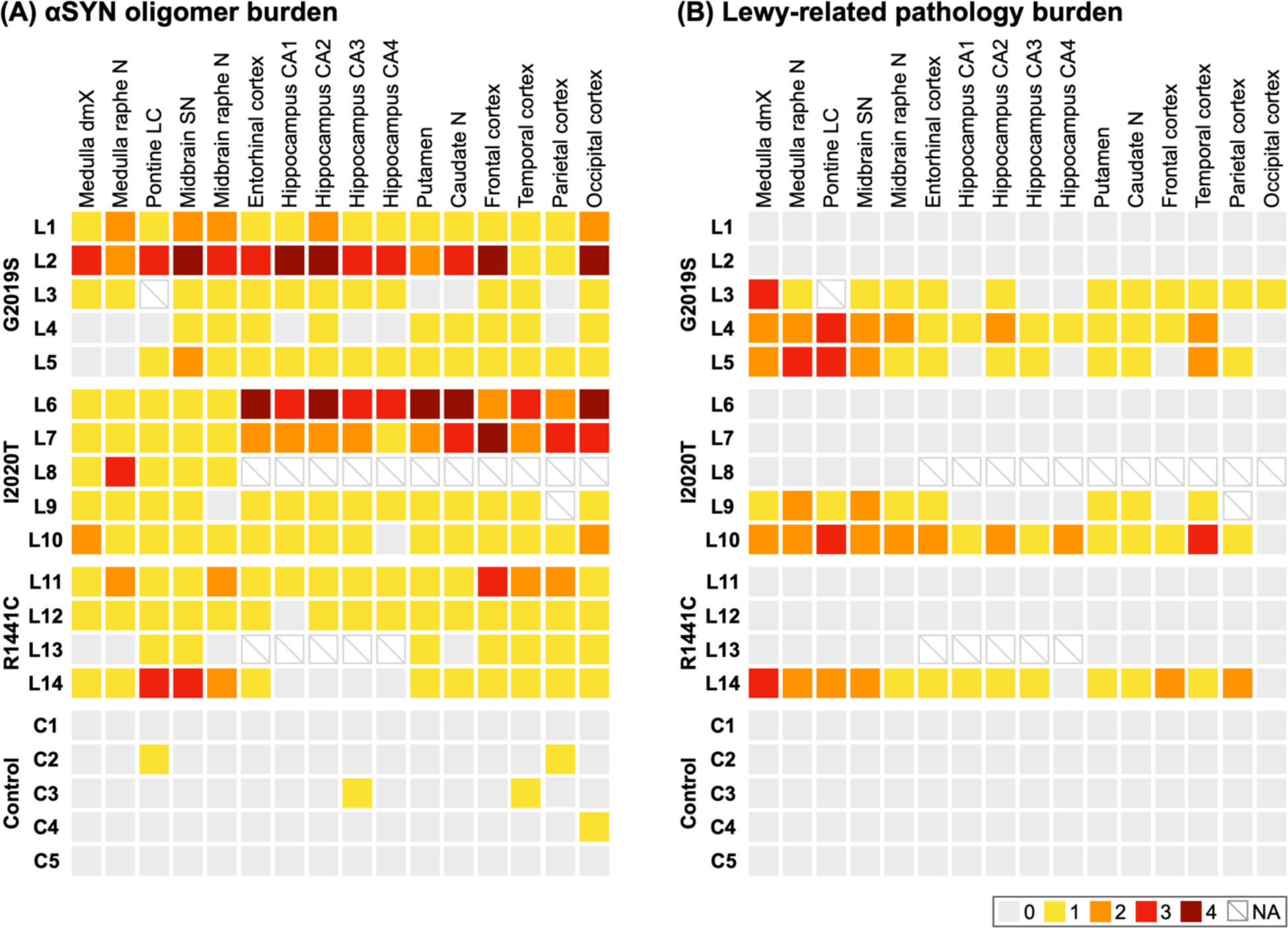
Comparison of α-synuclein oligomer burden (A) and Lewy-related pathology (B). *LRRK2*-PD patients without Lewy-related pathology demonstrate more prominent αSYN oligomer accumulation compared to those with Lewy-related pathology. *αSYN*, α-synuclein; *PLA*, proximity ligation assay; *dmX*, dorsal motor nucleus of the vagal nerve; *N*, nucleus; *LC*, locus coeruleus; *SN*, substantia nigra; *CA*, cornu ammonis.

### αSYN oligomers in each brain region

We compared the severity of αSYN oligomer burden for each *LRRK2* mutation to that of controls in each brain region (**Figure 4**). To mitigate the risk of type I errors associated with multiple comparisons, statistical analyses were confined to comparisons between individual mutation groups and the control group. LRRK2 p.G2019S exhibited a significantly greater αSYN oligomer burden in the substantia nigra, midbrain raphe nucleus, entorhinal cortex, hippocampal CA2 subfield, frontal cortex, and occipital cortex compared to controls. Similarly, p.I2020T showed significantly higher αSYN oligomer burden in the dorsal motor nucleus of the vagus nerve, medulla raphe nucleus, entorhinal cortex, and hippocampal CA1, CA2, and CA3 subfields, as well as in the putamen, caudate nucleus, and various cortical regions (frontal, temporal, parietal, and occipital) compared to controls. In contrast, p.R1441C demonstrated significant differences only in the substantia nigra and locus coeruleus compared to controls.

**Figure 4.**
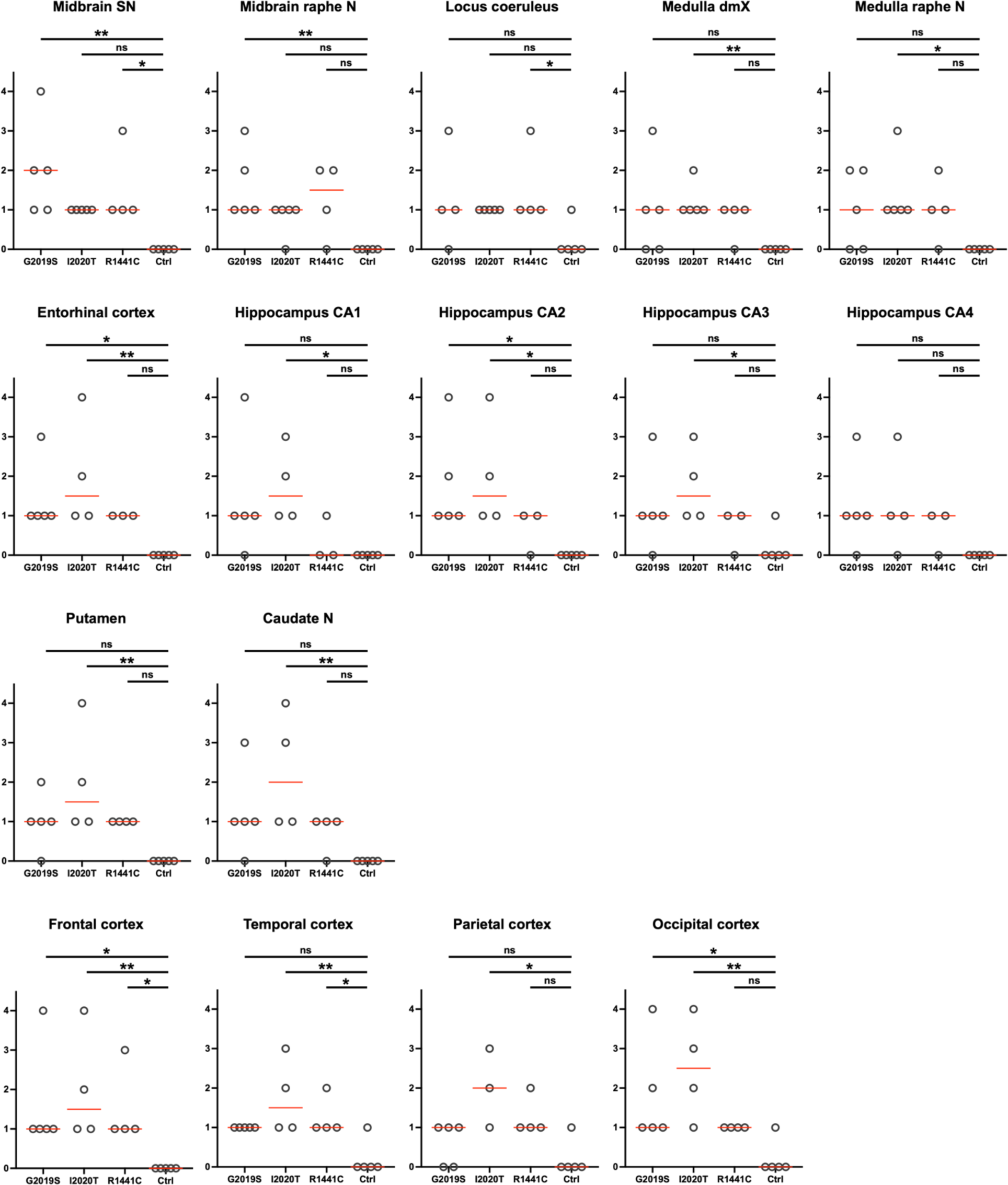
Severity of α-synuclein oligomer burden in each brain region of patients with *LRRK2* mutations and control subjects. *SN*, substantia nigra; *N*, nucleus; *dmX*, dorsal motor nucleus of the vagal nerve; *CA*, cornu ammonis.

### Correlation between clinical and pathological features

Next, we compared correlations between disease duration and pathological burden. The analysis revealed a trend for correlations between disease duration and pathological severity, but they did not reach statistical significance. The combined αSYN oligomer score showed a weak negative correlation with disease duration (r=-0.20 [-0.67, 0.39], P=0.49). This suggests a tendency for patients with shorter disease duration to have a higher burden of αSYN oligomers. In contrast, the Lewy-related pathology score demonstrated a weak positive correlation with disease duration (r=0.23 [-0.36, 0.69], P=0.42), indicating a slight tendency for increased Lewy-related pathology with longer disease duration, although this correlation was not statistically significant.

## Discussion

In the present study, we examined the distribution of αSYN oligomers in PD patients with three different *LRRK2* mutants: p.G2019S, p.I2020T, and p.R1441C. Approximately half of the patients did not have Lewy-related pathology; however, αSYN oligomers were detected in all patients regardless of the presence of Lewy-related pathology. Kalia, et al. reported that 65% (11/17) of p.G2019S, 11% (1/9) of p.I2020T, and 50% (2/4) of p.R1441C showed Lewy bodies [25]. Henderson, et al. reported that 56% (5/9) of p.G2019S patients exhibited Lewy-related pathology [21]. Clinical features of *LRRK2*-PD are largely indistinguishable from those of sporadic PD [20, 30]. Additionally, clinical features were comparable between *LRRK2*-PD patients with and without Lewy-related pathology [25]. To date, the accumulation of tau [4, 21, 48] or TDP-43 [2, 28, 40] has been reported as underlying pathology in a subset of *LRRK2*-PD; however, a unifying pathological change has not been identified. In this study, αSYN oligomers were detected in all cases, suggesting that αSYN oligomers may represent a common mechanism in *LRRK2*-PD. In general, parkinsonism due to tau or TDP-43 pathology does not respond to dopamine replacement therapy [7, 31]. In contrast, dopamine replacement therapy is effective in *LRRK2*-PD, which suggests that αSYN oligomers play a role in *LRRK2*-PD. The present study suggests that a therapeutic strategy for *LRRK2*-PD patients is targeting αSYN oligomers.

When comparing the severity of αSYN oligomer burden between cases with and without Lewy-related pathology, no significant difference was observed in p.R1441C. On the other hand, patients with p.G2019S and p.I2020T mutations without Lewy-related pathology had significantly greater αSYN oligomer burden compared to those with Lewy-related pathology. Additionally, a negative correlation was found between severity of Lewy-related pathology and αSYN oligomer burden. These findings suggest that αSYN oligomers are initially formed and accumulate in the brain; however, as Lewy-related pathology develops, the levels of αSYN oligomers may decrease. This observation aligns with previous findings on the temporal transition of αSYN aggregates in sporadic PD [44]. Specifically, in PD brains, αSYN oligomers accumulate first, followed later by Lewy-related pathology. As Lewy-related pathology develops, αSYN molecules are thought to accumulate within Lewy-related pathology, potentially providing a sink to decrease αSYN oligomers.

While αSYN oligomers were detected in all *LRRK2*-PD, Lewy-related pathology was observed only in about half of the patients. This observation suggests two possibilities. The first possibility is that in *LRRK2*-PD, the formation of Lewy-related pathology may take longer time compared to sporadic PD. This could explain why Lewy-related pathology is not observed in about half of *LRRK2* patients at autopsy [25]. The second possibility is that patients with a greater burden of αSYN oligomers may have a worse prognosis than those with fewer. Recent cellular and animal studies have demonstrated that αSYN oligomers are more toxic than Lewy-related pathology. Some studies have suggested a potential protective role of Lewy-related pathology against more toxic αSYN oligomers [38, 49, 51]. While speculative, high levels of αSYN oligomers may represent an aggressive outcome that shortens the lifespan in *LRRK2*-PD. Indeed, patients in the present study with the most abundant αSYN oligomers had a shorter disease duration.

To date, while no studies have examined αSYN oligomers in human brains with *LRRK2* mutations, several studies have investigated αSYN oligomers in cerebral spinal fluid (CSF) in subjects with *LRRK2* mutations. Aasly et al. investigated the levels of αSYN oligomers in the CSF of symptomatic and asymptomatic *LRRK2* mutation carriers and found that αSYN oligomers in the CSF were significantly elevated in asymptomatic *LRRK2* mutation carriers compared to healthy controls [1]. They also observed that CSF αSYN oligomer levels were inversely correlated with disease duration and disease severity in *LRRK2*-PD and sporadic PD patients. Another study by Majbour et al. also demonstrated higher levels of αSYN oligomers in the CSF in asymptomatic *LRRK2* mutation carriers and sporadic PD compared to healthy control [29]. The ratio of αSYN oligomers to total αSYN was higher in symptomatic and asymptomatic *LRRK2* mutation carriers and sporadic PD than in control subjects. In these studies, the relationship between αSYN oligomers in the CSF and αSYN oligomer accumulation in the brain remains unclear since correlative changes in the brains at autopsy were not available. Nevertheless, elevated levels of αSYN oligomers in the CSF in *LRRK2* mutation carriers suggest a possible link between *LRRK2* mutation and αSYN aggregation.

Some animal experiments have demonstrated an association between *LRRK2* and αSYN oligomers. Daher et al. showed that dopamine neuron loss, caused by the overexpression of αSYN or lipopolysaccharide exposure, was alleviated in *Lrrk2*-KO rats [10]. Another study using human p.A53T α-synuclein transgenic mice showed that α-synuclein pathology worsened when LRRK2 was overexpressed and that α-synuclein pathology was alleviated by LRRK2 ablation [27]. Ho et al. showed that LRRK2 inhibitor decreased αSYN oligomers in the striatum and phosphorylated αSYN in the cortex in p.R1441G mice [23]. Meanwhile, it should be noted that some studies have shown conflicting results [9, 22]. In light of these varying results, the present study, which revealed the accumulation of αSYN oligomers in postmortem human brains from *LRRK2*-PD patients, suggests a potential association between LRRK2 and αSYN oligomers. This finding highlights the importance of further investigating this relationship to better understand its implications in the pathogenesis of *LRRK2*-PD.

A limitation of the present study is the relatively small sample size. The number of patients for each *LRRK2* mutation ranged from 4 to 5, necessitating caution when interpreting the results due to the limited sample size. Nevertheless, this study comprehensively examined both early- and late-stage αSYN aggregates across multiple brain regions in patients with one of three different *LRRK2* mutations, thereby contributing new insights into the underlying pathology of *LRRK2*-PD.

## Conclusion

The present study revealed that αSYN oligomers were detected in all *LRRK2*-PD regardless of whether they had Lewy-related pathology at autopsy. Given the established toxicity of αSYN oligomers, their accumulation may play a crucial role in the pathogenesis of disease. These findings suggest that therapeutic strategies targeting αSYN oligomers may be warranted in *LRRK2*-PD.

## Acknowledgments

We sincerely gratitude to the patients and their families for their agreement to brain donation. We would like to acknowledge the continuous commitment, technical support and teamwork offered by Linda G. Rousseau, Whitney I. Davis, Virginia R. Phillips, and Monica Castanedes-Casey. This work was supported by the Mayo Clinic LBD Center WithOut Walls (CWOW; U54-NS110435), Ted Turner and family, the Little Family Foundation, and the Mayo Clinic Dorothy and Harry T. Mangurian Jr. Lewy body dementia program, and in part by the Mayo Clinic Florida Morris K. Udall Parkinson’s Disease Research Center of Excellence (NINDS P50 #NS072187), Alzheimer’s disease Research Center (P50 AG016574), an American Parkinson Disease Association (APDA) Mayo Clinic Information and Referral Center, an APDA Center for Advanced Research and the Mayo Clinic Lewy Body Dementia Association (LBDA) Research Center of Excellence.

## Conflict of interest

The authors declare no competing interests.

Ethical approval and consent to participate

All procedures performed in studies involving human participants were in accordance with the ethical standards of the institutional and/or national research committee and with the 1964 Helsinki Declaration and its later amendments or comparable ethical standards.

## Informed consent

Informed consent was obtained from the families of all individual participants included in the study.

## Author Contributions

H.S, T.T, Z.K.W, and D.W.D contributed to the study conception and design. Material preparation and data collection were performed by H.S, L.F, Y.H, M.T, K.N, N.F, K.H, Z.K.W, and D.W.D. H.S, K.H, H.K, O.A.R, P.J.M, T.T, Z.K.W, and D.W.D contributed to data analysis. H.S wrote the first draft of the manuscript and all authors commented on previous versions of the manuscript. All authors read and approved the final manuscript.

## Data Availability

The data that support the findings of this study are available from the corresponding author upon reasonable request.

---

**Supplementary Figure 1.**
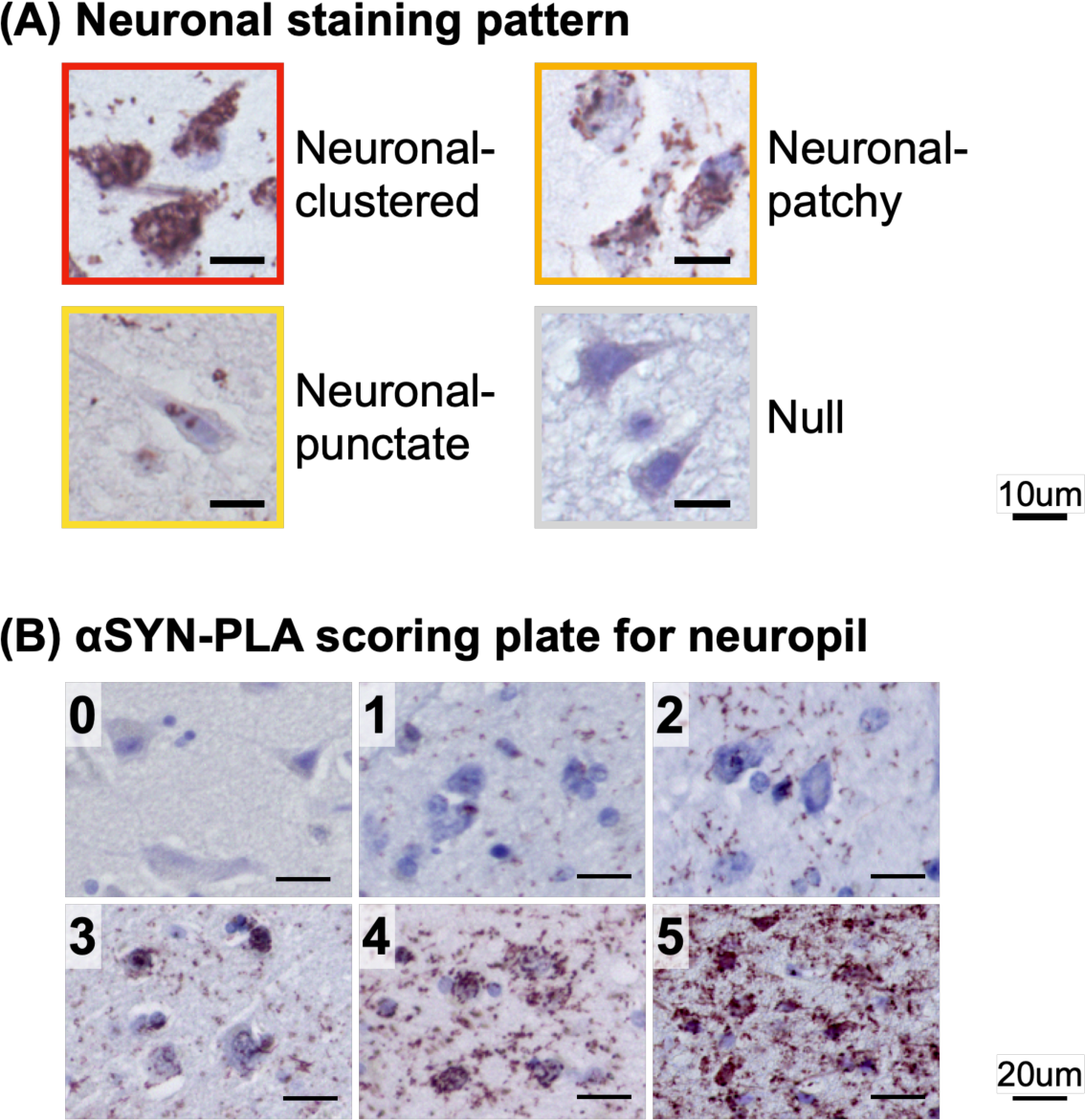
*αSYN*, α-synuclein: *PLA*, proximity ligation assay. (A) Neuronal staining pattern of αSYN-PLA was classified into 4 patterns: neuronal-clustered, neuronal-patchy, neuronal-punctate, and null. *Scale bars* 10 μm. (B) αSYN-PLA severity in neuropil was scored from 0 (no signal) to 5 (highest) by pre-made scoring plate. *Scale bars* 20 μm α*SYN*, α-synclein: PLA, proximity ligation assay

**Supplementary Figure 2.**
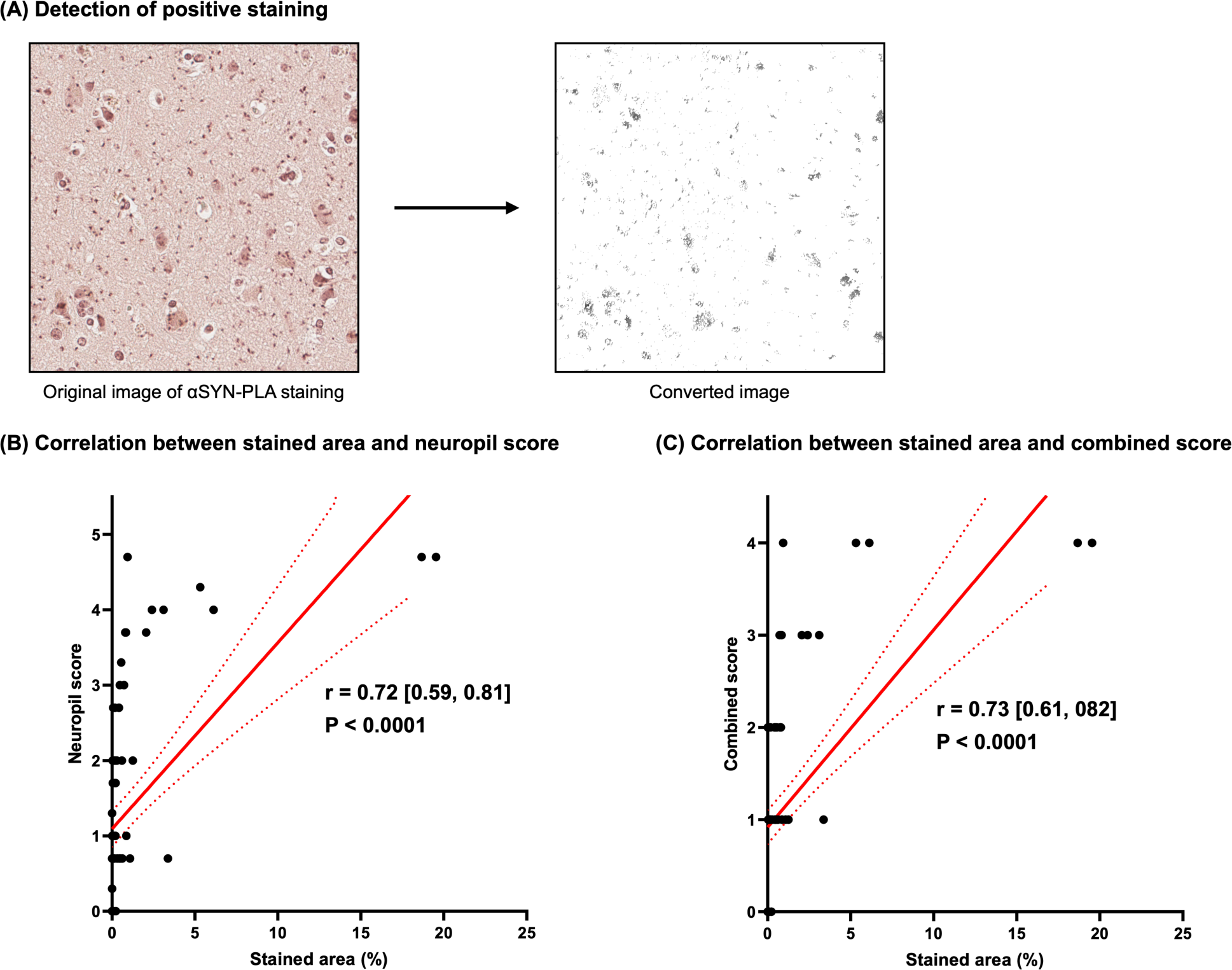
(A) Example of image conversion for stained area measurement of αSYN-PLA staining using the software ImageJ. (B) Correlation between stained area and neuropil score. (C) Correlation between stained area and combined αSYN oligomer score. *αSYN*, α-synuclein

**Supplementary Figure 3.**
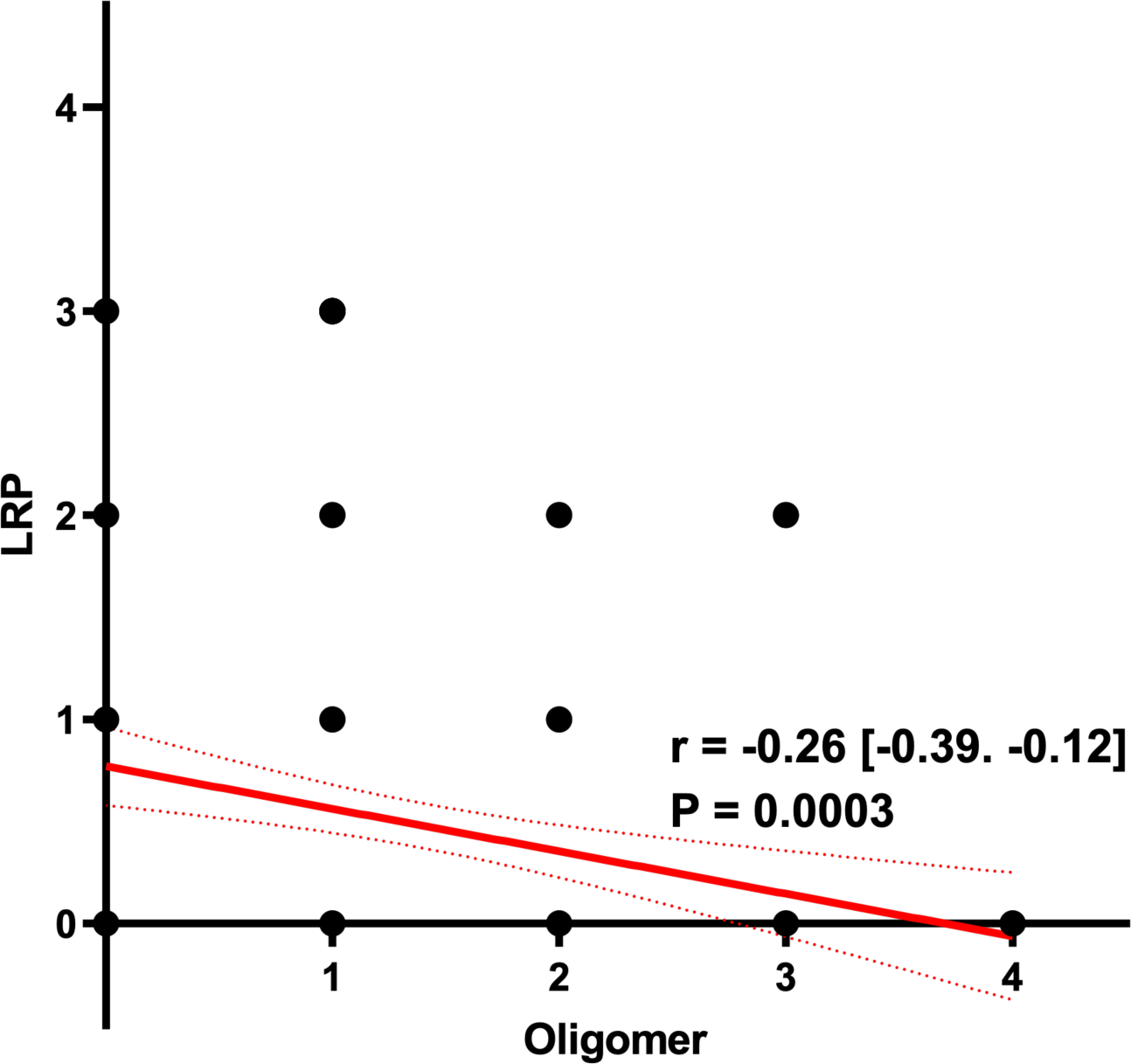
Correlation between αSYN oligomer burden and Lewy-related pathology burden. *αSYN*, α-synuclein: *LRP*, Lewy-related pathology.

